# Replication gaps are a cancer vulnerability counteracted by translesion synthesis

**DOI:** 10.1101/781997

**Authors:** Sumeet Nayak, Jennifer A. Calvo, Ke Cong, Emily Berthiaume, Jessica Jackson, Radha Charon Dash, Alessandro Vindigni, Kyle M. Hadden, Sharon B. Cantor

## Abstract

The replication stress response which serves as an anti-cancer barrier is activated not only by DNA damage and replication obstacles, but also oncogenes, mystifying how cancer evolves. Here, we identify that oncogene expression, similar to cancer therapies, induces single stranded DNA (ssDNA) gaps that reduce cell fitness, unless suppressed by translesion synthesis (TLS). DNA fiber analysis and electron microscopy reveal that TLS restricts replication fork slowing, reversal, and fork degradation without inducing replication fork gaps. Evidence that TLS gap suppression is fundamental to cancer, a small molecule inhibitor targeting the TLS factor, REV1, not only disrupts DNA replication and cancer cell fitness, but also synergizes with gap-inducing therapies. This work illuminates that gap suppression during replication is critical for cancer cell fitness and therefore a targetable vulnerability.

## INTRODUCTION

The replication stress response is activated in response to DNA lesions or intrinsic replication fork barriers, and it is critical to ensure the accurate transmission of genetic material to daughter cells (Berti and Vindigni, 2016). In response to stress, replication forks slow and remodel into reversed fork structures and this local fork response confers a signal to arrest DNA replication throughout the cell (Mutreja et al., 2018; Neelsen and Lopes, 2015). Cells either undergo replicative senescence or engage in DNA repair or other transactions that restart stalled DNA replication forks. The replication stress response is also induced by oncogenes, making it a critical barrier to cancer (Bartkova et al., 2005; Bartkova et al., 2006; Di Micco et al., 2006; Dominguez-Sola et al., 2007; Gorgoulis et al., 2005; Mailand et al., 2000; Neelsen et al., 2013; Yang et al., 2017). While oncogenes activate the DNA damage response, replication has been shown to accelerate and slow (Alevizopoulos et al., 1997; Neelsen et al., 2013; Santoni-Rugiu et al., 2000). These disparate findings could reflect distinct experimental systems or kinetics of analysis. Moreover, while checkpoint deficiency propels oncogenic transformation, (Bartkova et al., 2005; Bartkova et al., 2006; Di Micco et al., 2006), the oncogene-inducing lesion that limits fitness and is eventually overcome in cancer remains unknown. Here, we further queried how cellular replication responds to replication stress induced by drugs or oncogene expression.

A pathway known for tolerating DNA damage that interferes with replicative polymerases, is translesion synthesis (TLS). TLS polymerases are recruited to bypass replication blocking lesions when the replicative polymerases are not functional or physically blocked (Kannouche and Lehmann, 2004). TLS polymerases have been implicated in bypassing DNA damage induced by chemotherapies such cisplatin providing rational for TLS inhibition in cancer therapy (Doles et al., 2010; Wojtaszek et al., 2019; Xie et al., 2010b). However, we recently proposed a competing model in which single-stranded DNA (ssDNA) gaps underlie the mechanism-of-action of genotoxic therapies and that gap suppression is the key factor that confers resistance (Panzarino et al., 2019; Cong et al., 2019).

Here, we propose that the primary function of TLS is gap suppression (GS) to confer chemoresistance and overcome oncogene induced replication stress. Specifically, we show that TLS maintains continuous replication to limit ssDNA gaps induced by replication stress, oncogenes, or chemotherapy. We identify several cancer cell lines dependent on TLS for replication and fitness suggesting a TLS rewiring is essential for cancer initiation and/or evolution. Importantly, a small molecule inhibitor targeting the TLS factor, REV1, not only disrupts DNA replication and cancer cell fitness, but also synergizes with gap-inducing therapies. This work demonstrates that gap suppression is the fundamental mechanism of overcoming the anti-cancer barrier and that TLS inhibition is critical for therapy response.

## RESULTS

### TLS limits replication fork slowing during stress

To test the hypothesis that TLS avoids the replication stress response (**Figure 1A**), we sought to induce TLS and study its impact on the replication fork dynamics using DNA fiber spreading analysis. TLS is induced by the DNA helicase FANCJ when either its DNA damage induced acetylation or BRCA1 binding are blocked (Xie et al., 2010a; Xie et al., 2012). Here, we employed the FANCJ interaction defective BRCA1 mutant, FANCJ^S990A^, that promotes TLS via the polymerase polη (Xie et al., 2010a). We complemented FANCJ K/O U2OS, osteosarcoma cancer cells and FANCJ-null FA-J patient immortalized fibroblast cells, with FANCJ^S990A^ (pro-TLS), FANCJ^WT^ (control), or vector (V) and found as expected that both FANCJ^WT^ and FANCJ^S990A^ elevated mitomycin C (MMC) or cisplatin resistance as compared to vector (**Figure 1B, S1A and S1B**) (Peng et al., 2018; Xie et al., 2010a). In order to track the actively replicating fork, cells were labeled with sequential pulses of iodo-2′-deoxyuridine (IdU) and 5-chloro-2′-deoxyuridine (CldU) and the DNA tract lengths were measured. DNA fiber spreading analysis revealed that under unchallenged conditions, control or pro-TLS U2OS or FA-J complemented cell lines had similar tract lengths indicating that TLS induction did not impact normal replication progression. (**Figure S1C**). However, when CldU labeling was co-incident with 0.5mM hydroxyurea (HU), a dose that does not completely deplete nucleotide pools, but activates replication stress (Koc et al., 2004), or following ultraviolet radiation (UV), control cells had an expected reduction in tract lengths as compared to untreated control suggesting replication fork slowing during stress (**Figure 1C and S1D**). Notably, the DNA tracts in the pro-TLS cells failed to fully shorten during stress and appeared significantly longer than the control (**Figure 1C**). Further verifying that TLS contributes to the unrestrained replication, tracts fully shortened when the pro-TLS cells were treated with the TLS inhibitor (TLSi) that targets the C-terminus domain of REV (Korzhnev and Hadden, 2016; Sail et al., 2017) (**Figure 1C**).

**Figure 1.**
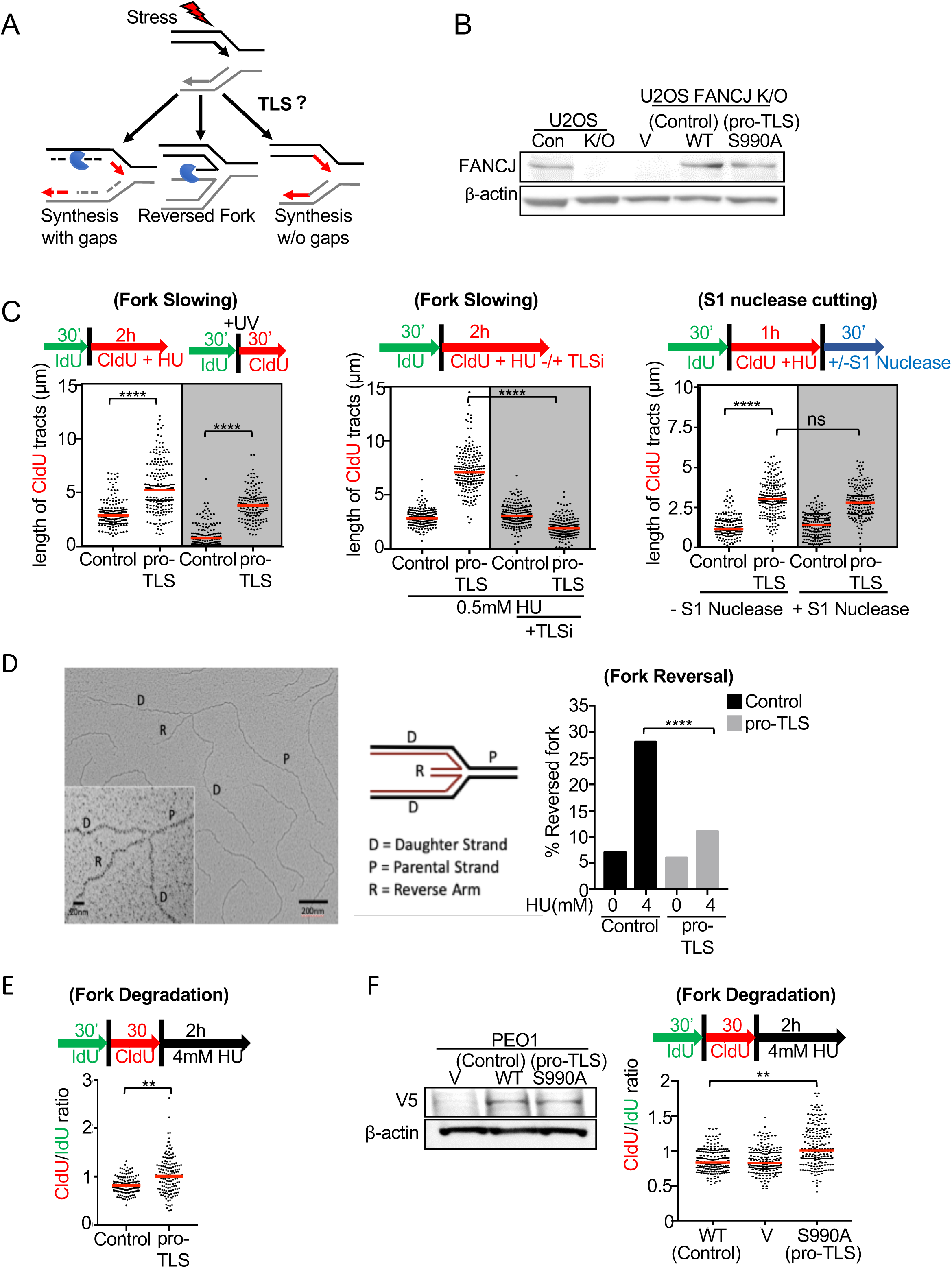
TLS limits replication fork slowing, reversal, degradation and gap induction. (A) Model to test whether TLS promotes unrestrained replication without gaps upon replication stress. Following DNA fiber analysis were performed in complemented U2OS FANCJ K/O cells. (B) Western blot analysis with the indicated Abs of WCE from U2OS CRISPR control and FANCJ CRISPR knockout (K/O) cells and FANCJ K/O cells complemented with empty vector (V), wild-type (WT) FANCJ and the FANCJ-BRCA1 binding deficient mutant (S990A). (C) Experimental schematic and quantification of CldU tract length after co-incubation with 0.5mM HU or 2J/m^2^ of UV or along with TLSi and quantification of CldU track length following −/+ S1 nuclease. (D) Quantification of the frequency of fork reversal following treatment with 4mM HU for 2h. Following number of replication intermediates were analyzed; untreated sample - 94 for (Control) and 84 for (pro-TLS), while for HU treated –130 for (Control) and 176 for (pro-TLS) from two independent experiments. (E-F) Experimental schematic and quantification of CldU/IdU ratio and western blot analysis with the indicated Abs of WCE from PEO1 cells expressing empty vector (V), FANCJ^WT^ (WT), and the FANCJ-BRCA1 binding deficient, FANCJ^S990A^ (pro-TLS). For every DNA fiber assay analysis, each dot represents one fiber. Experiments were performed in biological triplicate with at least 100 fibers per replicate. Bars represent the mean ± SD. Statistical analysis according to two-tailed Mann-Whitney test. All p values are described in Statistical methods.

One possibility for the longer tracts and the failure to slow replication in the pro-TLS cells could be a more rapid restart of stalled forks and/or the firing of new origins upon stress. However, this did not appear to be the case, based on the analysis of replication restart and new origin firing. In particular, after an initial IdU label, followed by a brief HU treatment, and final CldU label, control and pro-TLS cells readily incorporated CldU as compared to the vector complimented FANCJ null FA-J cells, that displayed replication restart defects and the aberrant activation of dormant origins (**Figure S1E**). Together, these findings suggest that TLS induction overcomes fork slowing, but does not alter restart or dormant origin firing in response to stress.

### TLS promotes replication fork progression during stress without ssDNA gap induction

Failure to slow replication during stress is associated with fork degradation, genomic instability, and low fitness (Lossaint et al., 2013; Neelsen and Lopes, 2015; Peng et al., 2018). We reasoned that TLS-dependent replication during stress could avoid this outcome because ssDNA gaps are avoided. To test this hypothesis, replication tracts were analyzed as before, but with or without S1 nuclease treatment. Indeed, we observed that pro-TLS cells generated significantly longer tracts that were maintained even after S1 nuclease treatment (**Figure 1C and S1F**). Given that S1 nuclease degrades DNA fibers with ssDNA gaps, within the labelled replication tracts, not observable in the standard DNA fiber assay (Quinet et al., 2017), these findings further indicate that the failure to slow is not associated with re-priming or new origin firing. In contrast, tract shortening with S1 nuclease occurred in FANCJ null cells or in cells expressing the FANCJ^K52R^ (helicase dead) or TLS inactivating mutant FANCJ^S990A+K52R^ (pro-TLS+ helicase dead) (Cantor et al., 2001; Xie et al., 2010a) (**Figure S1F**). Collectively these findings indicate that in response to stress, TLS not only disrupts fork slowing, but continues with the replication without generating ssDNA gaps.

### TLS avoids fork reversal and degradation

We predicted that continued replication during stress limited fork reversal and in turn protected replication forks from degradation. To investigate the frequency of reversed fork intermediates in cells with or without TLS induction, we analyzed the fine replication fork architecture by employing psoralen cross-linking coupled to Electron Microscopy (EM). Following 4mM HU treatment, we found a significant increase in the reversed fork structures in the control cells (∼28% reversed forks), whereas pro-TLS cells exhibited significantly lower frequency (∼11%) of fork reversal events (**Figure 1D**). Collectively, these results suggest that TLS restricts fork reversal.

Next, to asses fork degradation, we analyzed the ratio of CldU to IdU tract lengths following sequential pulses with IdU and CldU followed by HU treatment (Schlacher et al., 2011) (**Figure 1E**). Compared to control, the pro-TLS cells had a modestly enhanced CldU to IdU tract length ratio, consistent with less degradation upon stress (**Figure 1E and S1G**). Moreover, pro-TLS cells also maintained fork integrity following a prolonged period of replication stress (**Data not shown**). Additionally, in fork degradation-prone BRCA2-deficient PEO1 ovarian cancer cells(Sakai et al., 2009) (Kolinjivadi et al., 2017; Mijic et al., 2017; Schlacher et al., 2011; Taglialatela et al., 2017), ectopic expression of the pro-TLS mutant not only enhanced fork protection, but also conferred cisplatin resistance as forks failed to fully slow as compared to control (**Figure 1F and S1H**). Taken together, these findings indicate that TLS provides fork protection through suppression of fork remodeling similar to the loss of fork remodelers (Kolinjivadi et al., 2017; Peng et al., 2018; Vujanovic et al., 2017).

### TLS disrupts the global replication stress response without ssDNA induction

During stress, fork slowing and remodeling promote the global arrest of DNA replication and maintain genomic stability (Mutreja et al., 2018) (**Figure 1A**). Thus, TLS interference with fork slowing in response to stress could limit the global arrest of DNA replication. To test this idea, an asynchronous population of the control or the pro-TLS U2OS cell lines were either left untreated or treated with varying doses of HU while also being labeled with 5-ethynyl-2′-deoxyuridine (EdU) to track active replication. The efficiency of replication was quantified by scoring the number of EdU positive cells. Under unperturbed conditions, the EdU incorporation was similar between the control and pro-TLS U2OS cells, further suggesting that TLS does not impact the gobal replication in unchallenged conditions (**Figure 2A**). However, upon HU treatment, we observed that EdU incorporation was significantly reduced in the control U2OS cells, mimicking the replication fork slowing as studied by the DNA fiber assay (**Figure 2A**). Moreover, the pro-TLS U2OS cells, continued to incorporate EdU not only in HU, but also following treatment with ultraviolet radiation (UV) (**Figure S2A**) again suggesting that TLS promotes replication during stress.

**Figure 2.**
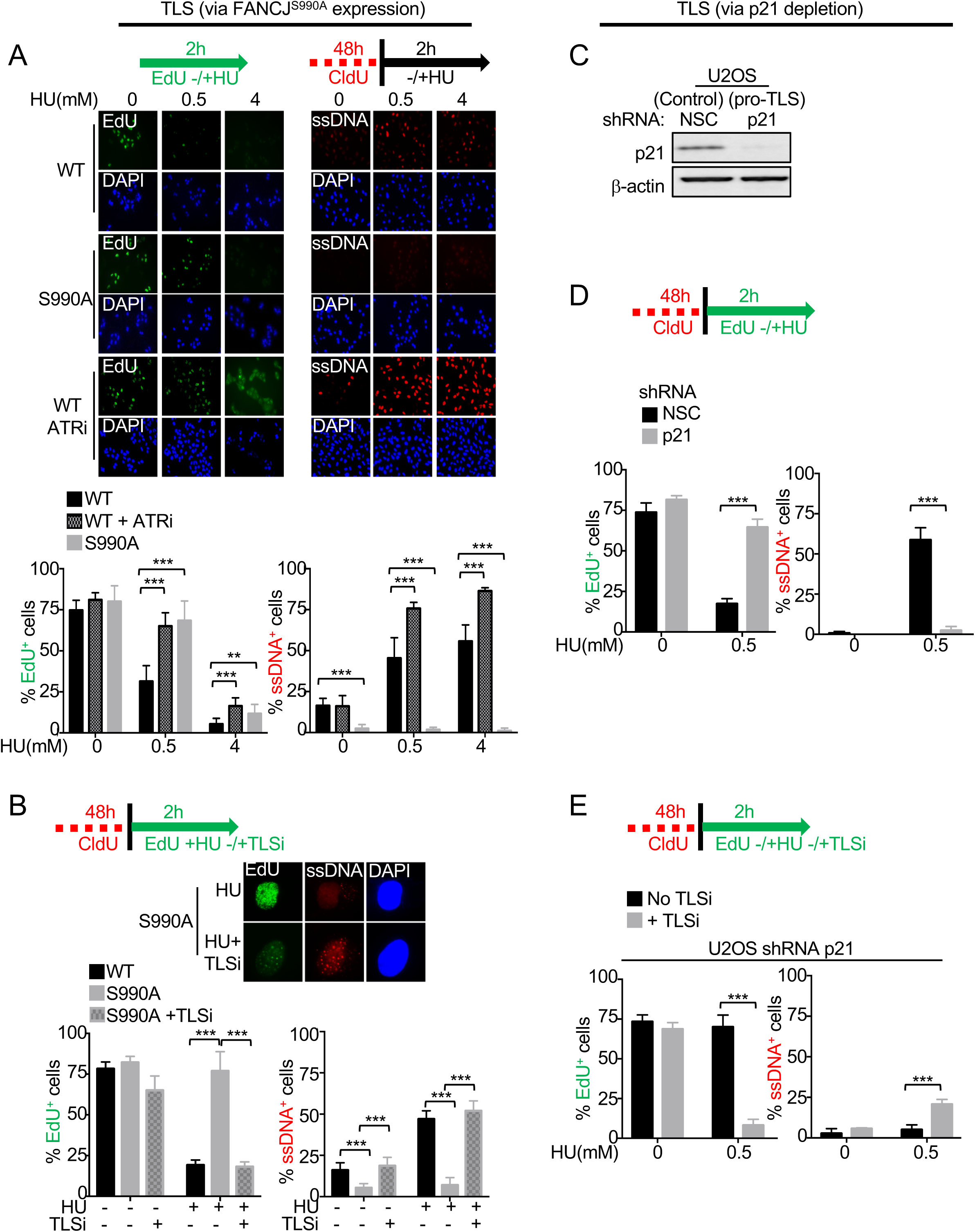
TLS promotes global replication during stress and suppresses ssDNA gaps. (A) Experimental schematic, representative immunofluorescence images and quantification of EdU and ssDNA positive cells. Cells were stained for EdU using the ClickiT chemistry and for ssDNA using CldU specific Ab under non-denaturing conditions and DAPI. Percent EdU and ssDNA positive cells were quantitated from around 300 cells counted from multiple fields. (B) Experimental schematic and representative immunofluorescence images at 100X following label with EdU in presence or absence of HU and/or TLSi and stained for EdU using the ClickiT chemistry and stained for ssDNA using ssDNA specific antibody and DAPI. (C) Western blot analysis with the indicated Abs of whole cell extract (WCE) from U2OS cells expressing shRNA against NSC or p21. (D-E) Experimental schematic and quantification of EdU and ssDNA positive cells. Cells were stained as described above. Experiments were performed in biological triplicate. Bars represent the mean ± SD. Statistical analysis according to two-tailed Mann-Whitney test. All p values are described in Statistical methods.

Inhibition of the checkpoint kinase ATR also enables replication during stress (Mutreja et al., 2018) (**Figure 2A**). Given that ATR inhibition is toxic to cells (Couch et al., 2013), we considered that a key difference between TLS activation and ATR inhibition was ssDNA gap induction. To test whether replication during stress differed by ssDNA gap induction, we performed non-denaturing immunofluorescence following incorporation of 5-chloro-2′-deoxyuridine (CldU). In the presence of HU, we observed that the ATRi lead to wide-spread global ssDNA gaps, whereas by comparison to control, the pro-TLS cells appeared resistant to gap formation (**Figure 2A**). Collectively, these findings indicate that TLS, unlike ATR inhibition, promotes replication during stress but without genome wide ssDNA gap induction.

To verify that unrestrained replication without ssDNA gaps is a distinct feature of TLS and not limited just to TLS-driven by FANCJ, we next depleted the negative regulator of TLS, p21 (Avkin et al., 2006). Depletion of p21 elevated continuous “ungapped” replication during HU treatment as compared to the control (**Figure 2C and S2B**). Importantly, in either pro-TLS system, FANCJ^S990A^ driven or on p21 depletion, the TLSi disrupted EdU incorporation and induced ssDNA gaps (**Figure 2B, 2E and S2C**). Notably, in unchallenged cells the TLSi did not interfere with EdU incorporation (**Figure 2B**) suggesting that the addition of stress was a pre-requisite to induce TLS-dependent replication. Collectively, these findings indicate that TLS is a robust mechanism for continuation of replication during stress without ssDNA gap formation.

### Oncogene expression induces ssDNA gaps and reduces cell fitness

Oncogene activation is associated with replication stress that serves as a barrier to cancer (Bartkova et al., 2005; Bartkova et al., 2006; Di Micco et al., 2006; Dominguez-Sola et al., 2007; Gorgoulis et al., 2005; Mailand et al., 2000; Neelsen et al., 2013; Yang et al., 2017). Given our findings, we sought to test the hypothesis that oncogene induced stress can be offset by TLS. To test this hypothesis, we generated cells stably infected with either empty vector -or *CCNE1* vector that encodes the oncogene cyclin E1 in a doxycycline inducible manner (DOX-ON system) (**Figure 3A**). As previously reported, we observed that cyclin E1 expression did not alter EdU incorporation (Alevizopoulos et al., 1997; Lukas et al., 1997; Santoni-Rugiu et al., 2000) (**Figure 3B, 3C, S3A and S3B**). However, there was a significant induction of genome wide ssDNA as well as loss of clonogenic capacity, both of which were suppressed by TLS maintained by expression of the FANCJ^S990A^ mutant (**Figure 3B, 3C, S3A and S3B**). Similar findings were observed in another well-established U2OS cyclin E1 inducible system (TET-OFF system) (Lukas et al., 1997). Upon cyclin E1 over-expression (OE), as compared to the normal ectopic levels (NE), EdU incorporation was maintained, but ssDNA was induced and clonogenic survival was reduced unless counteracted by TLS achieved by p21 depletion (**Figure 3D, 3E, 3F and S3C**). Furthermore, co-incubation of the TLSi restored cyclin E induced ssDNA gaps and the reduced clonogenic capacity of the pro-TLS cells (**Figure 3C and 3F**). Collectively these findings indicate that TLS buffers oncogene induced stress to promote “ungapped” replication and survival.

**Figure 3.**
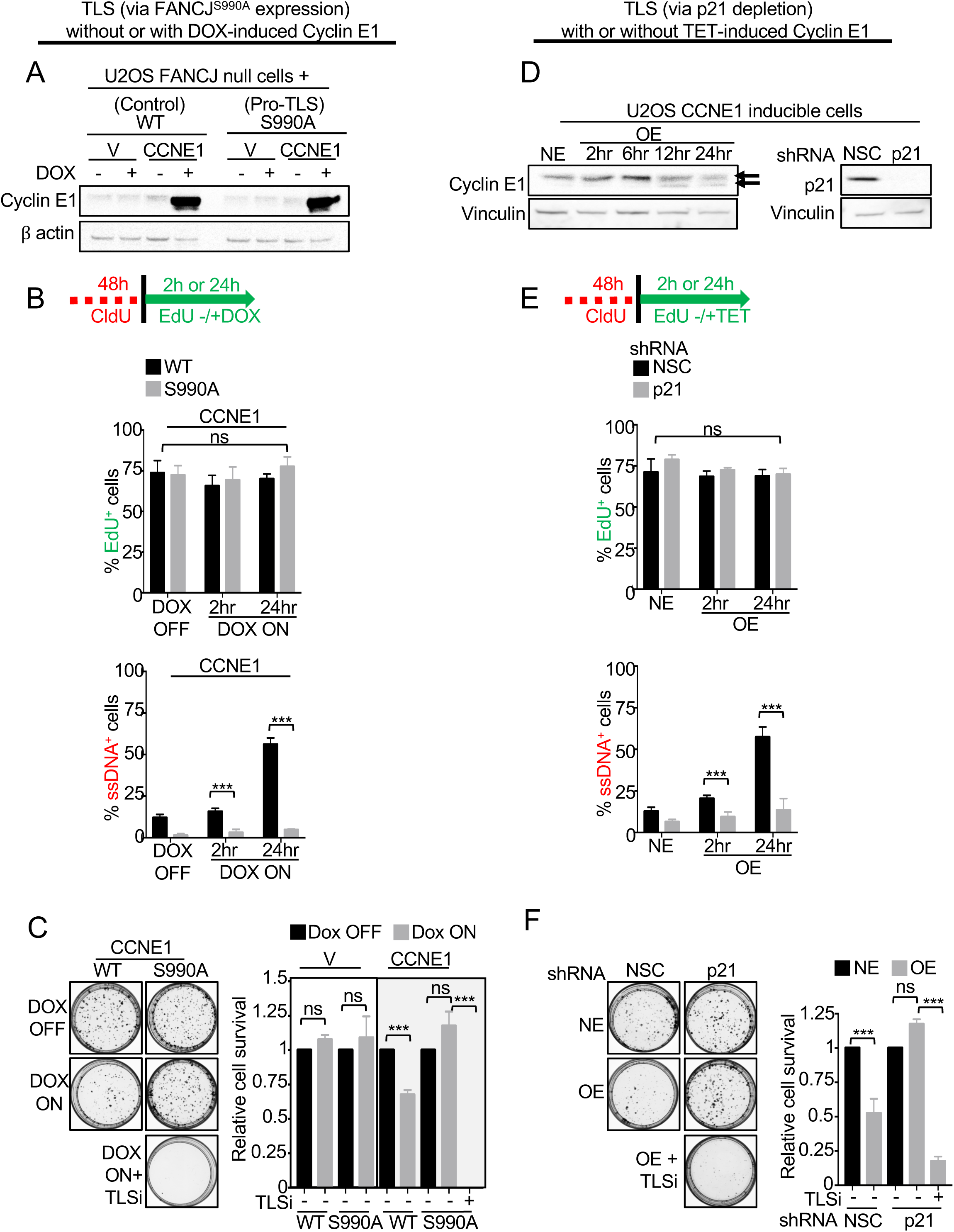
TLS overcomes oncogene induced stress response and promotes cell fitness. (A) Western blot analysis with the indicated Abs of WCE from FANCJ^WT^ or FANCJ^S990A^ complemented FANCJ K/O U2OS cell lines stably infected with either only vector (V) or vector with *CCNE1* gene. The cells were treated with 1ug/ml dose of doxycycline for 24hrs to induce cyclin E1 expression. (B and E) Experimental schematic and quantification of EdU and ssDNA positive cells. Cells were stained as previously described in Figure 2. (C) Representative images and quantification of the colony formation assay with and without doxycycline induced cyclin E1 expression. (D)Western blot analysis with the indicated Abs of WCE from U2OS cells inducibly expressing cyclin E1 and upon p21 depletion by using shRNA against NSC or p21. NE denotes normal level of cyclin E1 while OE denotes cyclin E1 overexpression. (F) Representative images and quantification of the colony formation assay in NSC vs p21 depleted U2OS cyclin E1 NE or OE cells. Experiments were performed in biological triplicate. Bars represent the mean ± SD. Statistical analysis according to two-tailed Mann-Whitney test. All p values are described in Statistical methods.

### Cancer cells show TLS dependence

If TLS overcomes the loss of fitness due to oncogene expression, then cancer evolution could favor TLS activation. To screen for a TLS rewiring in cancer, we tested the ability of distinct cancer cell types to replicate during stress. Remarkably, we found that replication robustly continued in the breast cancer cell line, MCF7, the endometrial cancer cell line, HeLa, the colon cancer cell line, HCT15, the lung cancer cell lines, A549 and NCI-H522, and the leukemia cell line, MOLT-4 (**Figure 4A, S4A and S4B**). Moreover, the TLSi curtailed replication during stress and induced ssDNA gaps in these cell lines (**Figure 4A and S4B**). Notably, MCF7 cells also showed a flattened morphology suggestive of senescence (**Figure 4A**). HeLa cells halted replication and induced ssDNA even in the absence of HU (**Figure 4A**), consistent with a pro-TLS phenotype even in unchallenged conditions. In contrast, similar to U2OS cells, the immortalized retinal pigment epithelial, RPE cell line ceased to replicate in low dose HU (**Figure S4A**).

**Figure 4:**
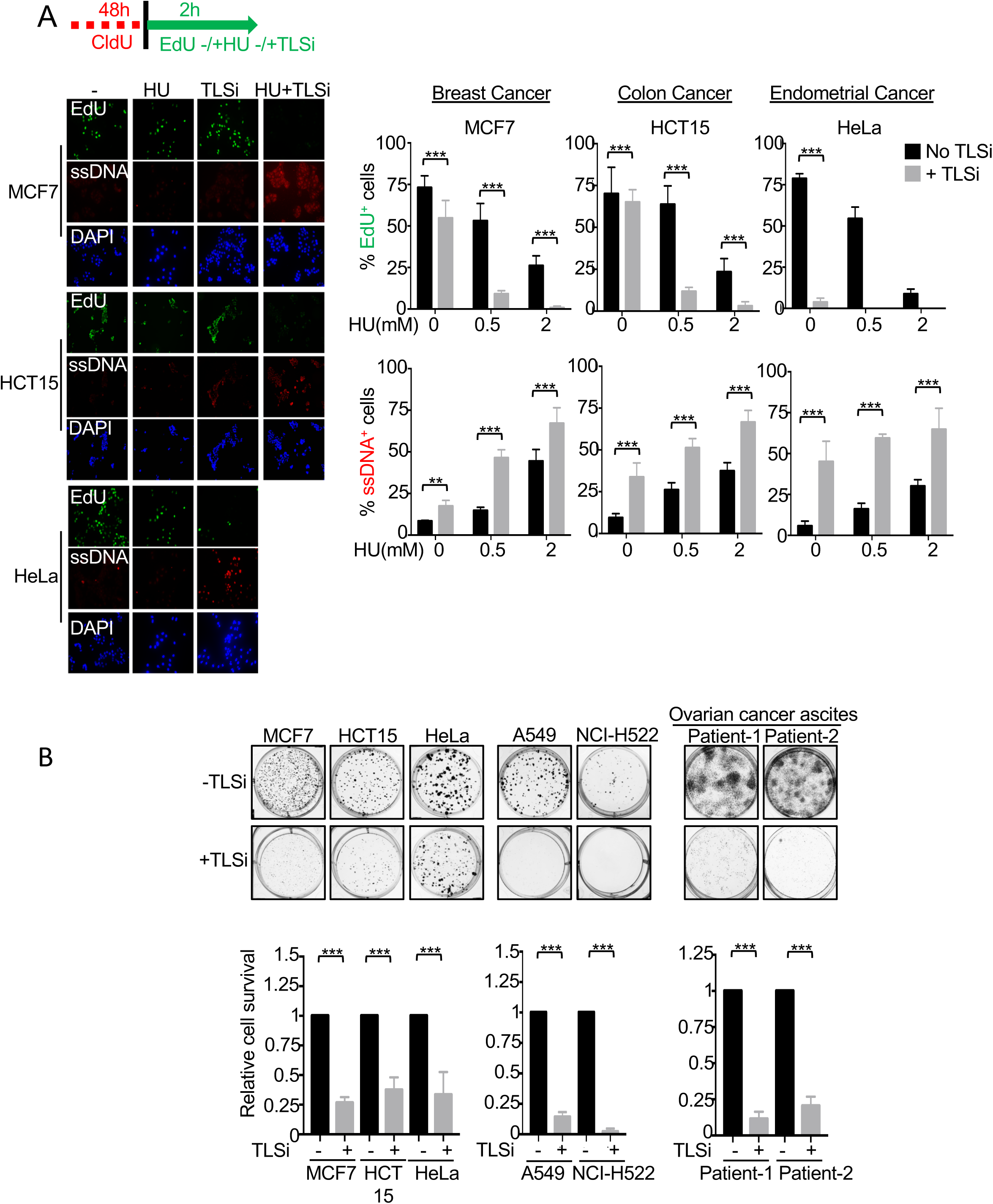
TLS subverts the replication stress response to promote cancer fitness. (A) Experimental schematic and quantification of EdU and ssDNA positive cells. Cells were stained as previously described in Figure 2. (B) Representative images and quantification of the colony formation assay with and without continuous presence of TLSi (20 μM). Experiments were performed in biological triplicate. Bars represent the mean ± SD. Statistical analysis according to two-tailed Mann-Whitney test. All p values are described in Statistical methods.

Consistent with a TLS rewiring, cancer cell lines with TLS-dependent replication lost clonogenic capacity upon treatment with the TLSi (**Figure 4B**). Moreover, early passage ovarian cancer ascites cells from two different patients were highly sensitive to the TLSi (**Figure 4B**). In contrast, the TLSi did not affect the colony forming capacity of the non-TLS dependent cell lines like RPE, U2OS and human mammary epithelial cell line, HMEC (**Figure S4C**). Additionally, TLS dependent Hela cancer cells showed dependence on the TLS factor FANCJ for replication and cellular fitness. Namely, FANCJ K/O in HeLa cells significantly reduced DNA replication and impaired clonogenic capacity (**Figure S5A and S5B**). Interestingly, along with the expected cisplatin sensitivity, p21 levels were also elevated in the FANCJ K/O HeLa cells and p21 depletion eventually improved replication, fitness and suppressed ssDNA gaps, unless TLS was inhibited (**Figure S5A, S5C and S5D**). Taken together, these findings reveal that distinct cancer cell lines rely on TLS for continuous replication and fitness indicating TLS is a cancer vulnerability.

### Gap inducing therapies are also evaded by TLS

Currently, there is a major clinical effort to combat cancer by induction of replication stress through inhibition of ATR or the mitotic checkpoint kinase, Wee1 (Bukhari et al., 2019). Given that these drugs induce replication gaps (**Figure 2A**) (Couch et al., 2013; Forment and O’Connor, 2018; Yang et al., 2017), we considered that if gaps were the sensitizing lesion, TLS could also interfere with their effectiveness. Indeed, compared to the non-TLS dependent cells, the pro-TLS U2OS cells had greater clonogenic survival following ATRi or Wee1i, inhibitor MK1775 (**Figure 5A, 5B and S6A-D**). Similarly, the pro-TLS cancer cell line, HCT15, also showed resistance to both the ATRi and Wee1i (**Figure 5A, 5B and S6A-D**) (Bukhari et al., 2019). However, when co-incubated with the TLSi, the pro-TLS U2OS or HCT15 cell lines were re-sensitized suggesting a more potent therapeutic response when TLSi is used in combination with ATRi or Wee1i (**Figure 5A, 5B and S6A-D**). Collectively, these findings demonstrate that TLS overcomes replication stress from oncogene expression that explains the prevalence of cancer cells rewired to depend on TLS for replication and fitness. This TLS rewiring mitigates the effectiveness of drugs such that inhibit ATR and Wee1, that induce gaps, suggesting the greater clinical potential of targeting TLS as a cancer therapy (**Figure 5C**).

**Figure 5:**
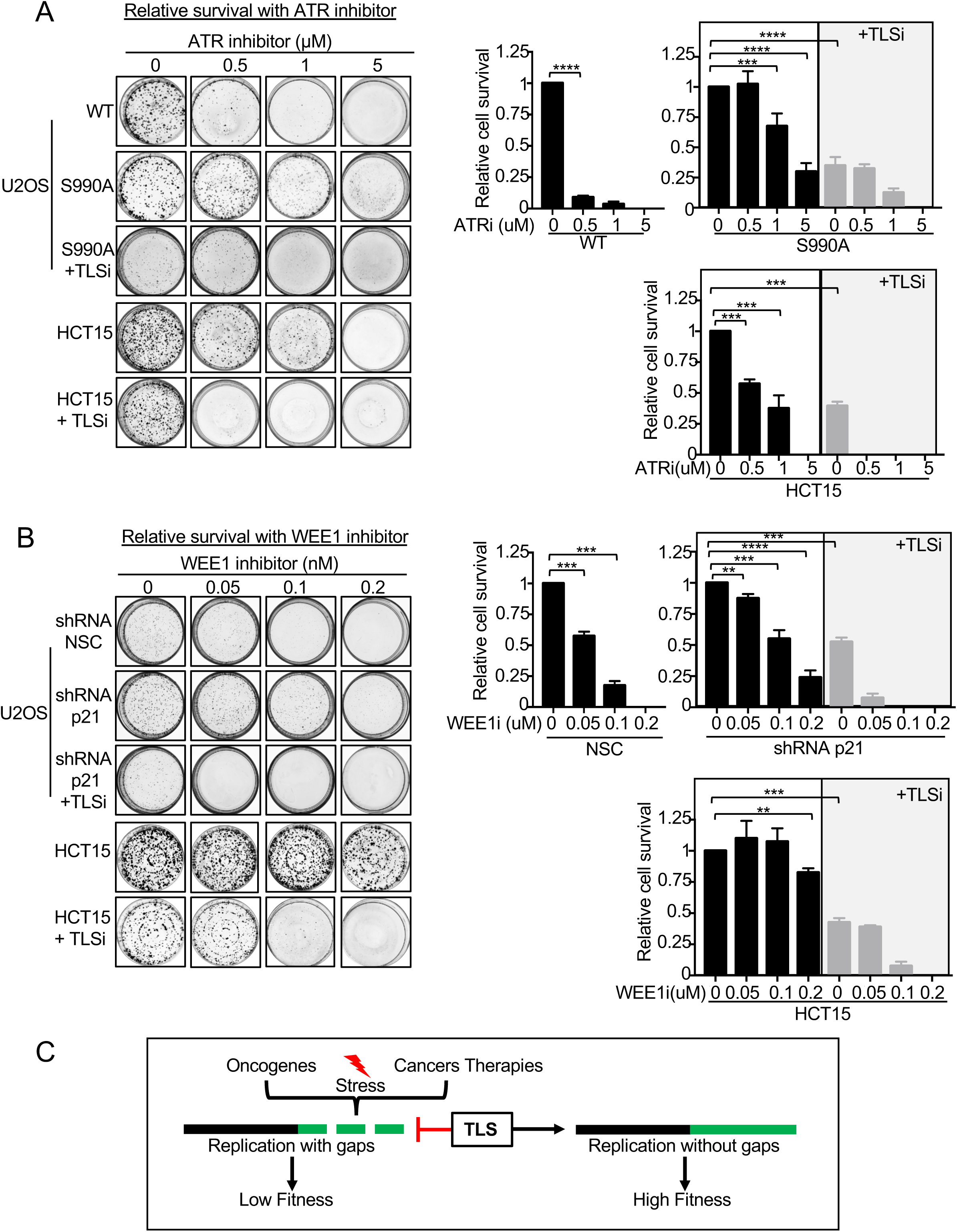
TLS as a gap suppression mechanism in cancer that can be re-sensitized by using the TLSi. (A-B) Quantification of the colony formation assay after dose dependent treatment with ATR and WEE1i alone or in combination with TLSi. (C) Model summarizing that TLS is a replication stress avoidance mechanism in cancer. Experiments were performed in biological triplicate. Bars represent the mean ± SD. Statistical analysis according to two-tailed Mann-Whitney test. All p values are described in Statistical methods.

## DISCUSSION

The replication stress response is a robust protective barrier that arrests or eradicates cells that are undergoing replication stress. Thus, cancers that overcome this barrier are expected to suppress the stress response by some mechanism. Based on our work, we predict that rewired replication that favors TLS is an essential adaptation to blunt oncogene induced replication stress that otherwise rapidly induces ssDNA gaps and limits fitness (**Figure 5C**). In support of this model, distinct cancer cells rely on TLS for fitness and to replicate during extrinsic or intrinsic stress. Mechanistically, we uncover that TLS curtails the slowing and remodeling of replication forks and the global replication arrest response while also suppressing ssDNA gaps that are toxic to cells (Chen et al., 1997; Forment and O’Connor, 2018; Gagna et al., 2000; Huang et al., 1996; Naruse et al., 1994; Nur et al., 2003; Peitsch et al., 1993; Tidd et al., 2000). Collectively, our findings highlight the importance of replication gaps as a cancer vulnerability that could be leveraged by TLS inhibition alone or in conjunction with gap inducing therapies in a wide range of cancers.

## Supplemental Figures

**Figure S1.**
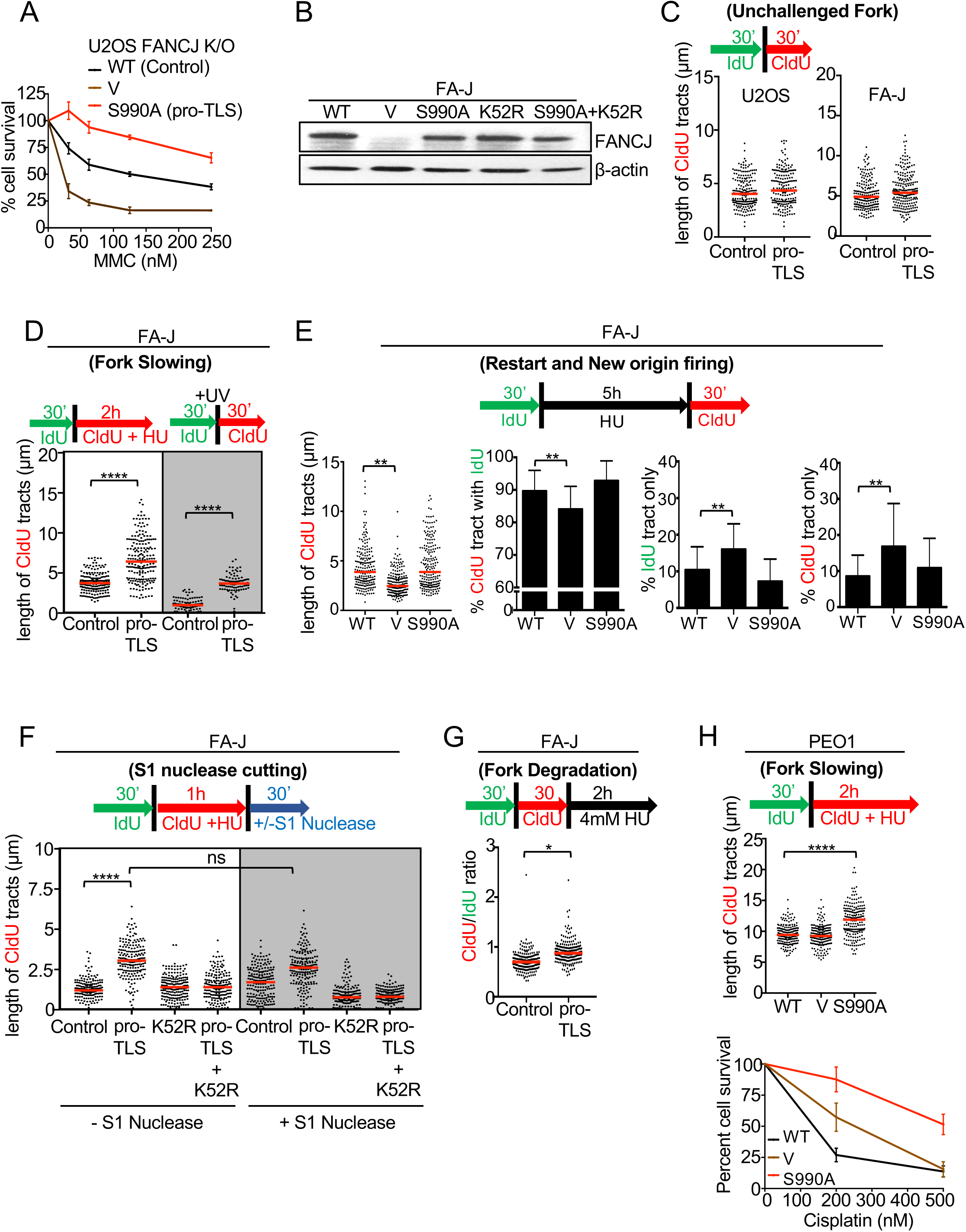
TLS does not impact replication restart or new origin firing. (A) Cell survival assays with U2OS FANCJ K/O complemented cells (complemented with FANCJ^WT^ (WT), empty vector (V) and the FANCJ-BRCA1 binding deficient, FANCJ^S990A^ (pro-TLS) treated with increasing concentrations of MMC. (B) Western blot analysis with the indicated Abs of lysates from FA-J cells complemented with FANCJ^WT^ (control), empty vector (V) and the FANCJ-BRCA1 binding deficient, FANCJ^S990A^ (pro-TLS), the FANCJ helicase dead mutant (K52R) and the FANCJ-BRCA1 binding deficient and helicase dead (K52R) double mutant. (C) Experimental schematic and the quantification of CldU tract length under unchallenged conditions as observed in the U2OS and the FA-J complemented control or pro-TLS cells. (D) Experimental schematic and the quantification of CldU tract length after co-incubation with 0.5mM HU or 2J/m^2^ of UV. (E) Experimental schematic and quantification of CldU tract length, percent CldU tract with IdU, percent only IdU and percent only CldU tract. (F) Experimental schematic and quantification of CldU track length following −/+ S1 nuclease. (G) Experimental schematic and quantification of CldU/IdU ratio. (H) Experimental schematic and quantification of CldU tract length. Cell survival assays with above three cell lines under increasing concentrations of cisplatin. Data represent the Mean percent ± s.d. of survival from three independent experiments. For every DNA fiber assay analysis, each dot represents one fiber. Experiments were performed in biological triplicate with at least 100 fibers per replicate. Bars represent the mean ± SD. Statistical analysis according to two-tailed Mann-Whitney test. All p values are described in Statistical methods.

**Figure S2.**
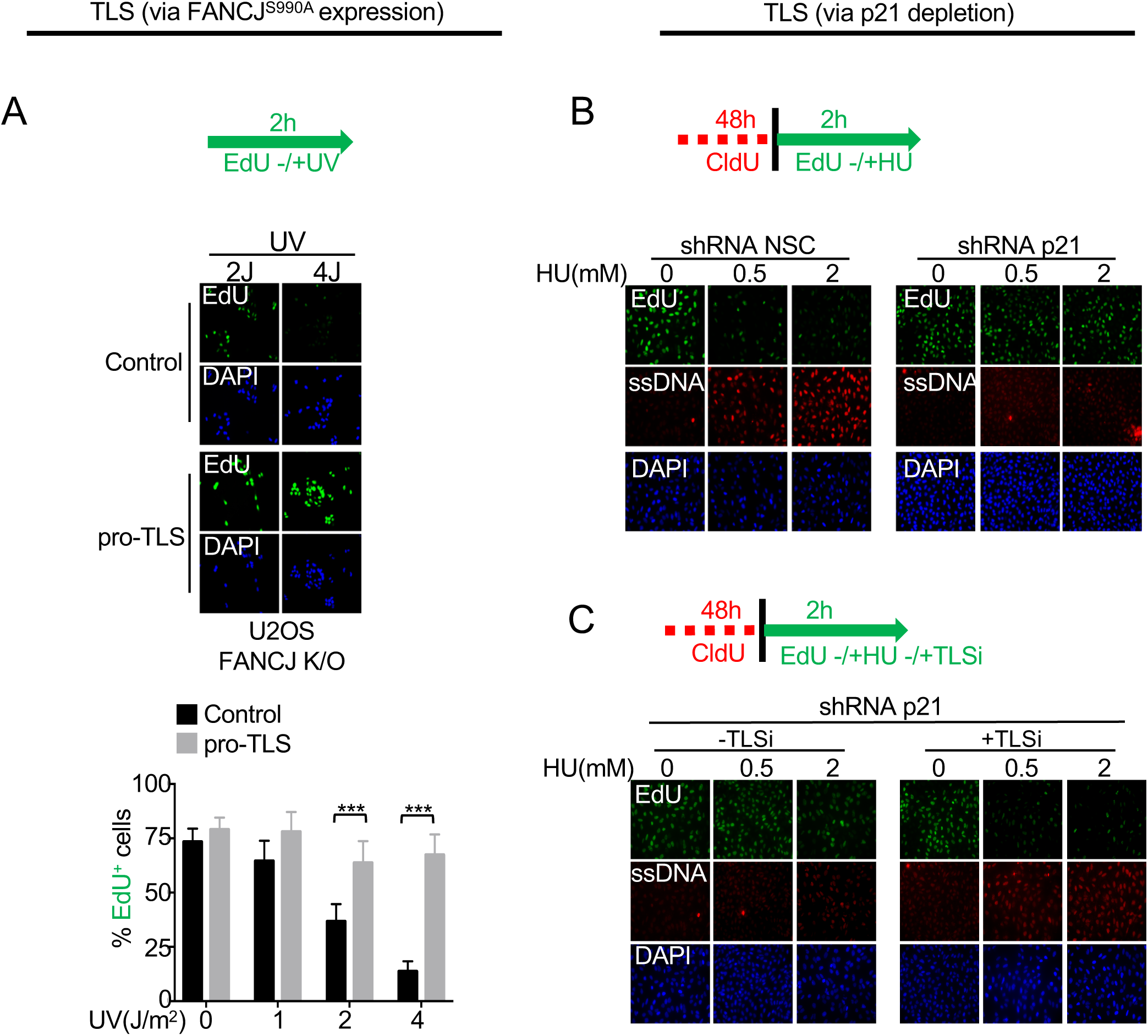
TLS overcomes replication stress associated ssDNA gaps. (A) Experimental schematic, representative images and quantification of the EdU positive cells after UV treatment. (B) Experimental schematic and representative images of EdU and ssDNA positive cells. Staining performed as described in main Figure 2. Experiments were performed in biological triplicate with at least 100 fibers per replicate. Bars represent the mean ± SD. Statistical analysis according to two-tailed Mann-Whitney test. All p values are described in Statistical methods.

**Figure S3.**
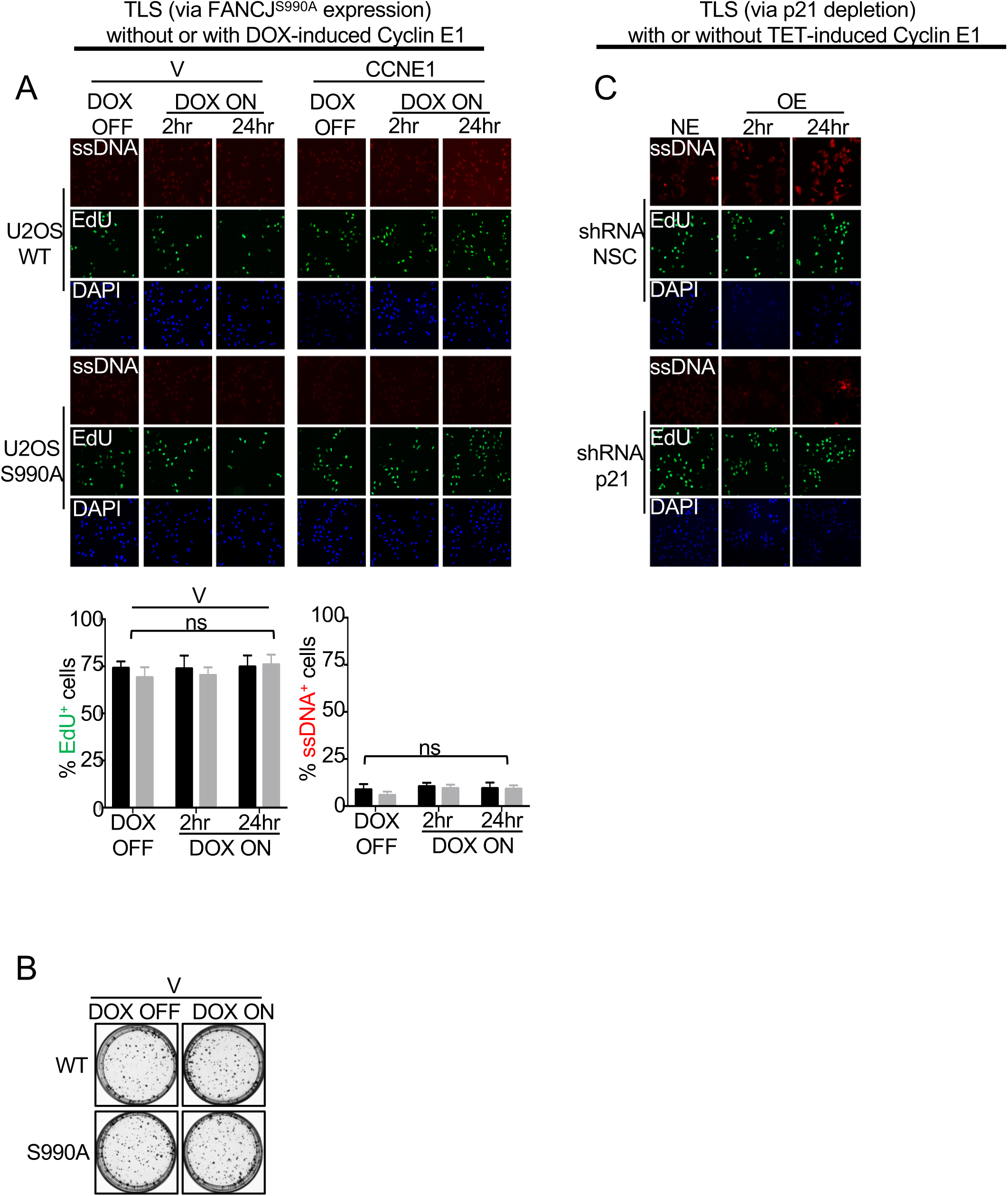
TLS counteracts oncogene induced replication stress response. (A and C) Experimental schematic, representative images and quantification of the EdU and ssDNA positive cells. Staining performed as described in main Figure 2. (B) Representative images of the colony formation assay upon doxycycline induction. Experiments were performed in biological triplicate with at least 100 fibers per replicate. Bars represent the mean ± SD. Statistical analysis according to two-tailed Mann-Whitney test. All p values are described in Statistical methods.

**Figure S4.**
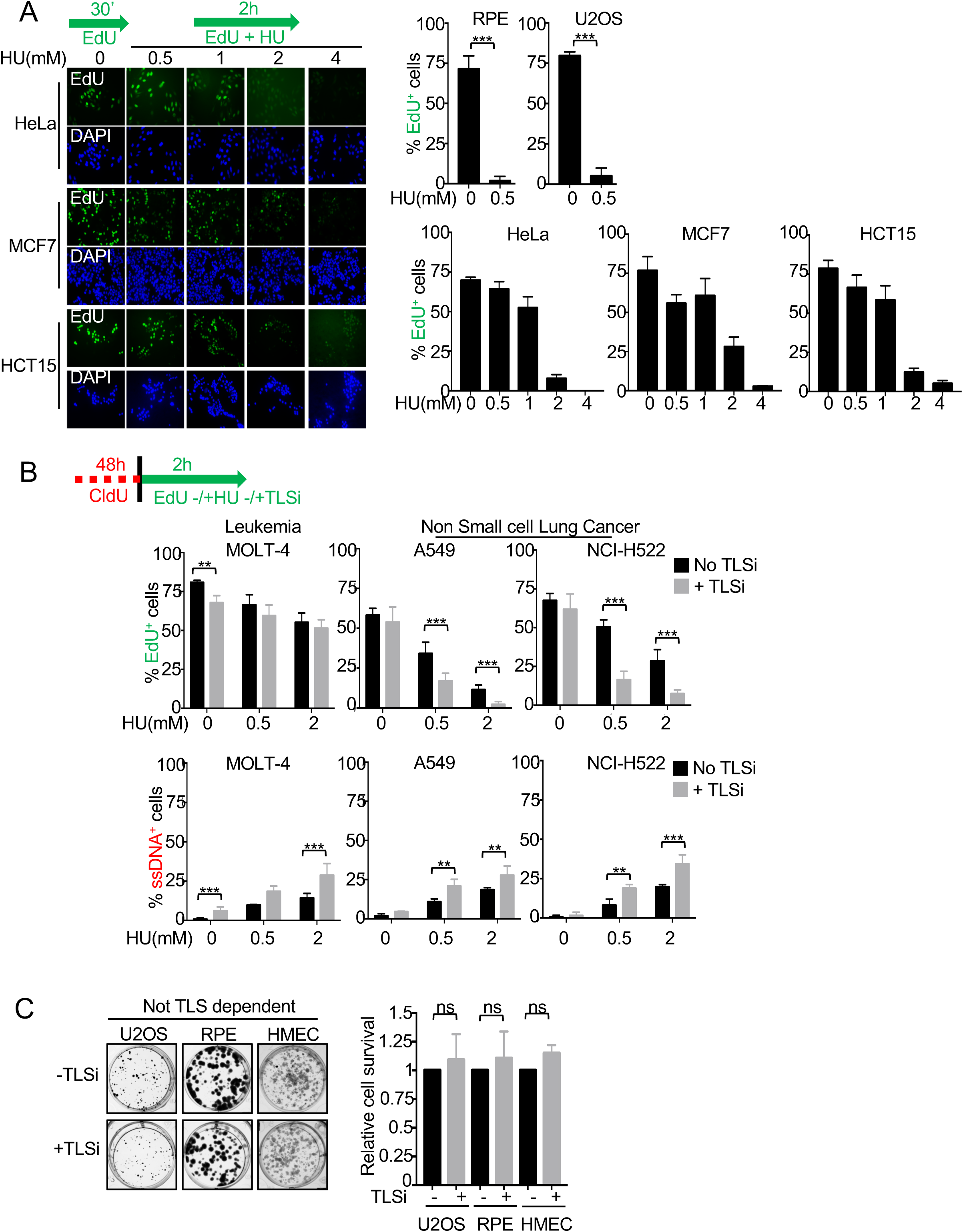
TLS re-wiring of cancer cells promotes unrestrained replication during stress. (A) Experimental schematic and quantification of EdU positive cells. Cells were stained for EdU using the ClickiT chemistry and DAPI. Percent EdU positive cells were quantitated from around 300 cells counted from multiple fields. (B) Experimental schematic and representative immunofluorescence images of EdU and ssDNA positive cells. Cells were stained as described previously in main figure 2. (C) Representative images of the colony formation assay with and without continuous presence of TLSi (20 μM). Experiments were performed in biological triplicate with at least 100 fibers per replicate. Bars represent the mean ± SD. Statistical analysis according to two-tailed Mann-Whitney test. All p values are described in Statistical methods.

**Figure S5.**
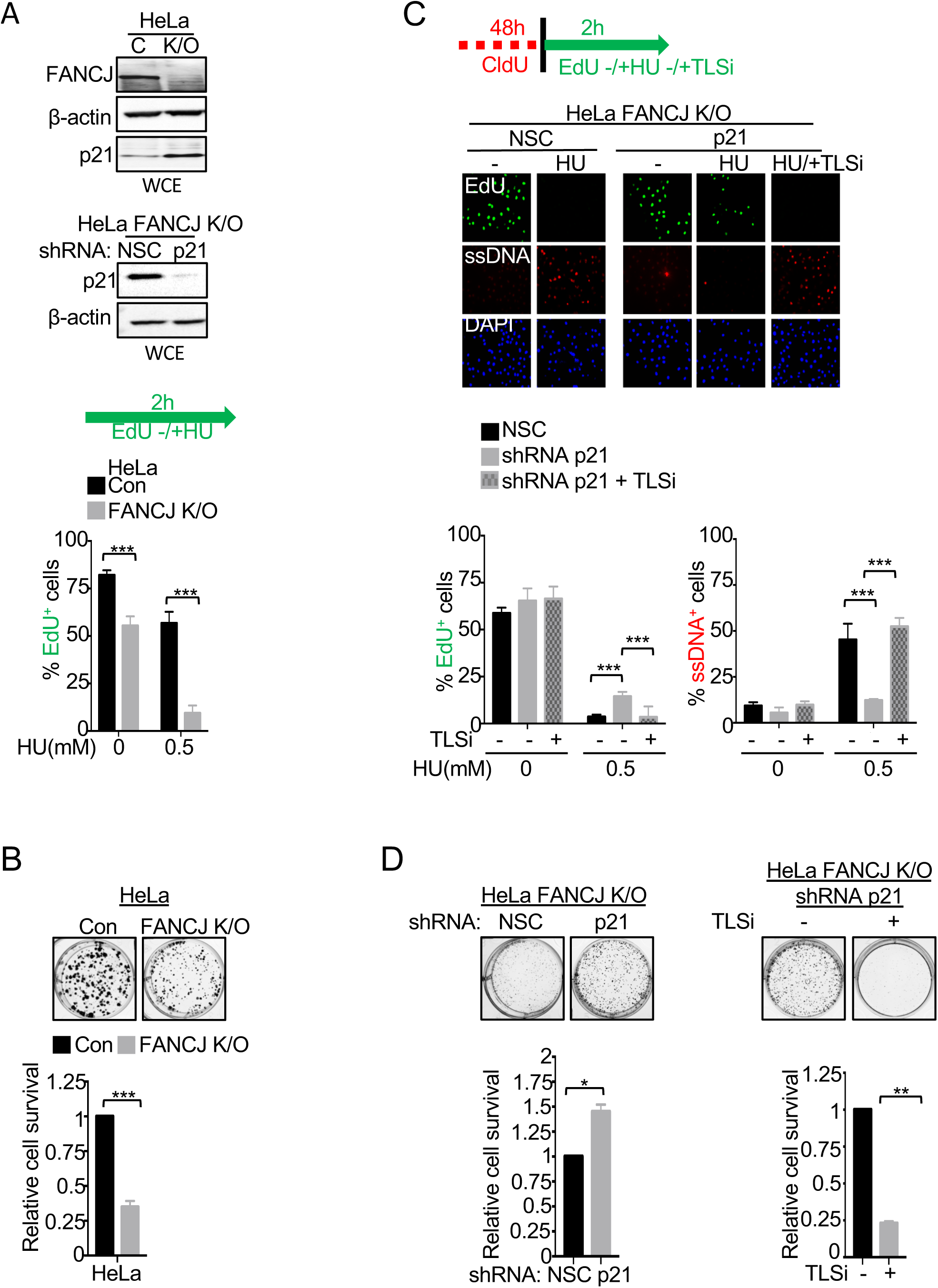
FANCJ promotes TLS by suppressing the negative regulator, p21. (A) Western blot analysis with the indicated Abs of whole cell extract (WCE) from HeLa CRISPR control and FANCJ CRISPR knockout (K/O) cells and HeLa FANCJ K/O cells expressing shRNA against NSC or p21. Experimental schematic and quantification of EdU positive cells. Cells were stained as previously described in main Figure 2. (B-D) Representative images and quantification of the colony formation assay. (C) Experimental schematic and quantification of EdU and ssDNA positive cells performed as described in main Figure 2. Experiments were performed in biological triplicate with at least 100 fibers per replicate. Bars represent the mean ± SD. Statistical analysis according to two-tailed Mann-Whitney test. All p values are described in Statistical methods.

**Figure S6.**
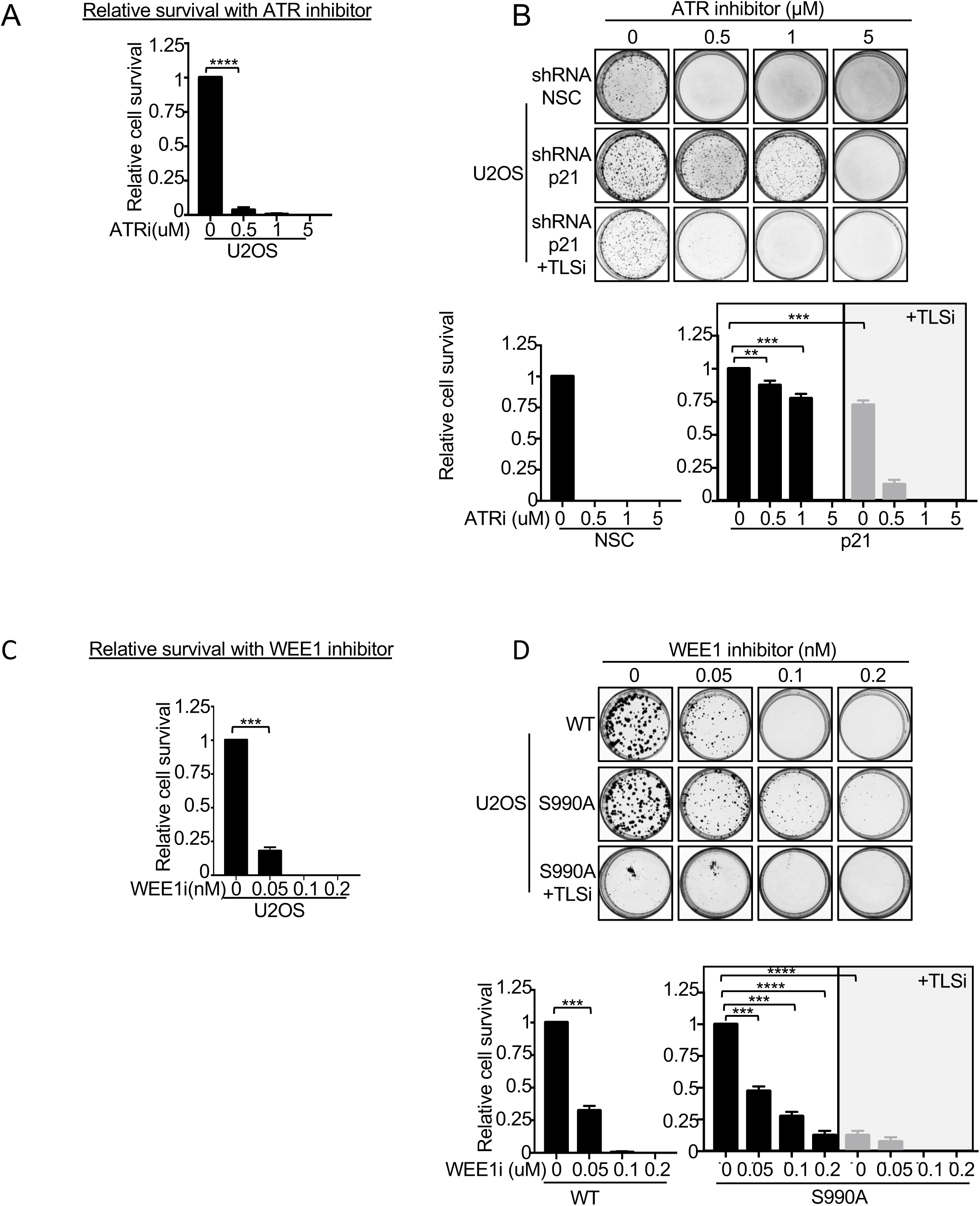
TLSi as a promising cancer therapy. (A-D) Representative images of the colony formation assay after dose dependent treatment with ATR and WEE1i alone or in combination with TLSi.

## EXPERIMENTAL PROCEDURES

### Cell lines

U2OS, PEO1, HeLa, MCF7, HCT15 and A549 cell lines were grown in DMEM supplemented with 10% fetal bovine serum and penicillin and streptomycin (100 U/ml each). U2OS cells inducibly overexpressing cyclin E (U2OS-CE) were maintained in DMEM supplemented with 10% fetal bovine serum (FBS; Invitrogen, Cat. No. 10500), penicillin-streptomycin-glutamine (Invitrogen, Cat. No. 10378-016), G418 400 µg/ml (Invitrogen, Cat. No. 10131-027), puromycin 1 µg/ml (Sigma, Cat. No. P8833) and tetracycline 2 µg/ml (Sigma, Cat. No. T7660). Right before the experiment, the cells were split into two aliquots. One aliquot was cultured in media without tetracycline to induce cyclin E overexpression (OE cells) and the other in media with 1µg/ml doxycycline to maintain low levels of ectopic cyclin E expression (NE cells). MOLT4 and NCI-H522 cell lines were grown in RPMI supplemented with 10% fetal bovine serum and penicillin and streptomycin (100 U/ml each). HMEC cell line was grown in DMEM supplemented with 15% fetal bovine serum and penicillin and streptomycin (100 U/ml each). FA-J cells (EUFA30-F) were immortalized with human telomerase reverse transcriptase (hTERT) (Peng et al., 2007) and cultured as previously described (Achar et al., 2015; Litman et al., 2005). Stable FA-J pOZ complemented cell lines were generated and selected as previously described (Peng et al., 2007). U2OS and HeLa FANCJ K/O CRISPR cell lines were generated as previously described (Peng et al., 2018). Stable U2OS FANCJ K/O and PEO1 pLenti complemented cell lines were generated by blasticidin selection (5 μg/ml). Stable HeLa and U2OS shRNA knockdown cell lines were generated by puromycin selection (0.25 and 0.5ug/ml respectively).

### Human Subjects

Malignant ovarian cancer cells were recovered from ascitic fluids from ovarian cancer patients by the University of Massachusetts Medical School (UMMS) Biorepository and Tissue Bank. Patient consent was obtained prior to specimen collection under an UMMS Institutional Review Board (IRB)-approved protocol (H4721). Malignant ovarian cancer cells were recovered from ascitic fluids by centrifugation at 200 g and cryopreserved in RPMI media supplemented with 10% FBS and 10% DMSO. Cells were slowly frozen at −80° C in an isopropanol bath overnight and stored long term in the vapor phase of a liquid nitrogen freezer. The consent process included conditions for sharing de-identified samples and information with other investigators. No identifiable information will be shared at any time per Health Insurance Portability and Accountability Act guidelines.

### Plasmid and shRNA Constructs

The WT and S990A FANCJ pLentiviral vectors were a gift from J Chen (Yu et al., 2003). The WT and S990A pOZ vectors were generated as previously described (Peng et al., 2007). HeLa FANCJ K/O and U2OS cells were infected with pLK0.1 vector containing shRNAs against non-silencing control (NSC) or one of three shRNAs against p21/CDKN1A (A) (Target region – 3’UTR - CGCTCTACATCTTCTGCCTTA), (B) (CDS - GAGCGATGGAACTTCGACTTT) and (C) (CDS -GTCACTGTCTTGTACCCTTGT). pInducer20 empty vector and pInducer20 Cyclin E1 plasmids were obtained from Addgene. The FANCJ K/O U2OS cells complemented with FANCJ WT or FANCJS990A were further infected with the respected virus to express the empty vector or Cyclin E1 in a doxycycline inducible manner. shRNAs were obtained from UMMS shRNA core facility.

### Drugs and reagents

The following drugs were used in the course of this study: Cisplatin (Sigma-Aldrich), was prepared as a 1mM solution in saline per the manufacturer’s instructions. MMC (Sigma-Aldrich), was prepared by dissolving 0.5mg/mL in water. HU (Sigma), was prepared fresh in complete media prior to experiments per the manufacturer’s instructions. The ATR inhibitor, VE-821 (Selleckchem) and WEE1i, MK-1775 (Selleckchem) were prepared as a 15mM and 5mM solutions in DMSO respectively. 5-chloro-2′-deoxyuridine (CldU) and 5-Iodo-2′-deoxyuridine (IdU) were obtained from Sigma-Aldrich. Click-iT EdU Alexa Fluor 488 Imaging Kit was obtained from Invitrogen. Concentration and duration of treatment are indicated in the corresponding figures and sections.

### Immunoblotting and Antibodies

Cells were harvested, lysed and processed for Western blot analysis as described previously using 150mM NETN lysis buffer (20mM Tris (pH 8.0), 150mM NaCl, 1mM EDTA, 0.5% NP-40, 1mM phenylmethylsulfonyl fluoride, 10mg/ml leupeptin, 10mg/ml aprotinin). For cell fractionation, we isolated cytoplasmic and soluble nuclear fractions with the NE-PER Kit (Thermo) according to the manufacturer’s protocol; to isolate the chromatin fraction, the insoluble pellet was resuspended in RIPA buffer and sonicated in a BioRuptor according to the manufacturer’s protocol (Medium power, 20 minutes, 30 seconds on, 30 seconds off at 4°C). Proteins were separated using SDS–PAGE and electro-transferred to nitrocellulose membranes. Membranes were blocked in 5% not fat dry milk(NFDM) phos-phate-buffered saline (PBS)/Tween and incubated with primary Ab for overnight at 4°C. Antibodies (Abs) for Western blot analysis included anti-βactin (Sigma-Aldrich), anti-FANCJ (E67), anti-Vinculin (Sigma-Aldrich), anti-Cyclin E1 (abcam) and anti-p21 (BD Pharmingen),. Membranes were washed, incubated with horseradish peroxidase-linked secondary Abs (Amersham) for 1h at room temperature and detected by chemiluminescence (Amersham).

### Immunofluorescence

Immunofluorescene was performed as described previously (Cantor et al., 2001). Cells were grown on coverslips in 10 µM BrdU for 48 h before the treatment with drugs. Cells were then treated with the aforementioned drugs for 2h. After treatment, cells were washed with PBS and pre-extracted (0.5% Triton X-100 made in phosphate-buffered saline (PBS) on ice. Cells were then fixed using 4% Formalin for 15 min at RT. Fixed cells were then incubated with primary Abs against BrdU (Abcam) at 37°C for 1h. Cells were washed and incubated with secondary Abs for 1 h at room temperature. After washing, cover slips were mounted onto glass slides using Vectashield mounting medium containing DAPI (Vector Laboratories). For EdU labeling, staining was carried out with Click-iT EdU imaging kit (Invitrogen) according to the manufacturer’s instructions.

### DNA Fiber Assays

To directly visualize replication fork dynamics, we established single molecular DNA fiber analysis. In this assay, progressing replication forks in cells were labeled by sequential incorporation of two different nucleotide analogues, 5-Iodo-2’-deoxyuridine (IdU, 50μM) and 5-Chloro-2’-deoxyuridine (CldU, 50μM), into nascent DNA strands for the indicated time and conditions. After nucleotide analogues were incorporated *in vivo*, the cells were collected, washed, spotted (2.5μl of 10^5^ cells/ml PBS cell suspension) and lysed on positively charged microscope slides (Globe Scientific, #1358W) by 7.5μl spreading buffer (0.5%SDS, 200mM Tris-Hcl pH 7.4, 50mM EDTA) for 8 min at room temperature. Individual DNA fibers were released and spread by tilting the slides at a 45 degree. After air dry, the fibers were fixed by 3:1 methanol: acetic acid at room temperature for 3 min. After air dry again, fibers were rehydrated in PBS, denatured with 2.5M HCl for 30 min, washed with PBS and blocked with blocking buffer (3% BSA and 0.1%Trition in PBS) for 1 hour. Next, slides were incubated for 2.5h with primary Abs for (IdU: 1:100, mouse monoclonal anti-BrdU, Becton Dickinson 347580; CldU: 1:100, rat monoclonal anti-BrdU, Abcam 6326) diluted in blocking buffer, washed several times in PBS, and then incubated with secondary Abs (IdU: 1:200, goat anti-mouse, Alexa 488; CldU: 1:200, goat anti-rat, Alexa 594) in blocking buffer for 1h. After washing and air dry, the slides were mounted with Prolong (Invitrogen, P36930). Finally, the visualization of green and/or red signals by fluorescence microscopy (Axioplan 2 imaging, Zeiss) will provide information about the active replication directionality at single molecular level.

### S1 Nuclease Fiber Assay

As described previously, cells were exposed to 50uM IdU to label replication forks, followed by 50uM CldU with 0.5mM HU for 1h. Subsequently, cells were permeabilized with CSK buffer (100 mM NaCl, 10 mM MOPS, 3 mM MgCl2 pH 7.2, 300 mM sucrose, 0.5% Triton X-100) at room temperature for 8 minutes, followed by S1 nuclease (20U/ml) in S1 buffer (30 mM Sodium Acetate pH 4.6, 10 mM Zinc Acetate, 5% Glycerol, 50 mM NaCl) for 30 minutes at 37C. Finally, cells were collected by scraping, pelleted, resuspended in 100-500ul PBS; 2ul of cell suspension was spotted on a positively charged slide and lysed and processed as described in the DNA fiber assay section above

### Electron microscopy

For the EM analysis of replication intermediates, 5–10 × 10^6^ U2OS cells treated with or without 4mM HU were collected and genomic DNA was cross-linked by two rounds of incubation in 10 μg ml^−1^ 4,5′,8-trimethylpsoralen (Sigma-Aldrich) and 3 min of irradiation with 366 nm UV light on a precooled metal block^10,26^. Cells were lysed and genomic DNA was isolated from the nuclei by proteinase K (Roche) digestion and phenol-chloroform extraction. DNA was purified by isopropanol precipitation, digested with PvuII HF in the proper buffer for 3–5 h at 37 °C and replication intermediates were enriched on a benzoylated naphthoylated DEAE–cellulose (Sigma-Aldrich) column. EM samples were prepared by spreading the DNA on carbon-coated grids in the presence of benzyl-dimethyl-alkylammonium chloride and visualized by platinum rotary shadowing. Images were acquired on a transmission electron microscope (JEOL 1200 EX) with side-mounted camera (AMTXR41 supported by AMT software v601) and analyzed with ImageJ (National Institutes of Health). EM analysis allows distinguishing duplex DNA—which is expected to appear as a 10 nm thick fiber, after the platinum/carbon coating step necessary for EM visualization—from ssDNA, which has a reduced thickness of 5–7 nm. The criteria used for the unequivocal assignment of reversed forks include the presence of a rhomboid structure at the junction itself in order to provide a clear indication that the junction is opened up and that the four-way junction structure is not simply the result of the occasional crossing of two DNA molecules^25^. In addition, the length of the two arms corresponding to the newly replicated duplex should be equal (b = c), whereas the length of the parental arm and the regressed arm can vary (a ≠ b = c ≠ d). Conversely, canonical Holliday junction structures will be characterized by arms of equal length two by two (a = b, c = d).

### Viability assays

Cells were seeded onto 96 well plates (500 cells per well, performed in triplicates for each experiment) and incubated overnight. Next day the cells were treated with increasing dose of MMC for 1 hr in serum-free media or cisplatin or TLSi and maintained in complete media for 5 days. Percent survival was measured using CellTitre-Glo viability assay (Promega) photometrically in a microplate reader (Beckman Coulter – DTX 880 Multimode detector).

### Colony Formation Assay

For colony formation assays either 500 or 1000 cells per well were seeded into 6 well plates and were treated continuously with or without different drugs as mentioned in the respective figures. Once the colonies had developed, the cells were fixed with 90% methanol and then stained with 0.05% (w/v) crystal violet solution. Plates were then imaged using ChemiDoc^TM^ Touch Imaging System (BioRad) and number of colonies were counted using the Cell Profiler software version 3.1.5 from Broad Institute.

### Statistical Methods

Statistical differences in DNA fiber assay, S1 Nuclease assay, Immunofluorescence and colony forming assays were determined using a two-tailed Mann-Whitney test. Statistical analysis was performed using Excel and GraphPad Prism (Version 7.0). In all cases, ns: not significant (p > 0.01), **p < 0.01, *p< 0.001, and ****p < 0.0001.

## ACKNOWLEDGMENTS

We thank the members of the Cantor laboratory for helpful discussions. We thank Dr. Michael Green for providing HCT15, A549, MOLT-4 and the NCI-H522 cell lines, Dr Thanos D. Halazonetis for providing the U2OS cyclin E1 inducible cell line. The work in the S.C. laboratory was supported by R01 CA176166-01A1, R01 CA225018-01A1, and the Basser Center for BRCA research, as well as charitable contributions from Mr. and Mrs. Edward T. Vitone, Jr. and the Lipp Family Foundation. The work in the A.V. laboratory is supported by NIH grant R01GM108648 and by DOD BRCP Breakthrough Award BC151728. The work in KH laboratory is supported by University of Connecticut Research Foundation. We also thank Dr. Karl Simin and the University of Massachusetts Medical School (UMMS) Biorepository and Tissue Bank for providing the human patient ovarian cancer ascites sample. The University of Massachusetts Medical School (UMMS) Biorepository and Tissue Bank is supported by the National Center for Advancing Translational Sciences of the National Institutes of Health under award number UL1-TR001453.

## Author contributions

S.B.C., S.N., K.H., and A.V. designed the experiments. S.N., J.A.C., K.C., E.B., and J.J. performed the experiments. S.N. and J.A.C. analyzed the data. R.C.D. generated the reagents. S.B.C. and S.N. wrote the manuscript. S.B.C., K.H. and A.V. supervised the research.

## DECLARATION OF INTERESTS

The authors declare no competing interests.

## Notes

The authors declare no potential conflicts of interest

